# A Bayesian approach to volumetric analysis in rare neurodegenerative disorders

**DOI:** 10.64898/2026.10.02.756349

**Authors:** Sharon Yuen Shan Ho, Tatjana Schmidt, Boyd Ghosh, James B. Rowe, Timothy Rittman

## Abstract

Identifying regional brain atrophy is critical for understanding neurodegenerative disorders. Traditionally, group comparisons rely on frequentist statistics, utilising *p*-values and point estimates that offer limited characterization of uncertainty. Bayesian methods provide an alternative by estimating effect sizes and credible intervals while incorporating prior knowledge and uncertainty. This study evaluated the feasibility of Bayesian hierarchical modelling to assess regional brain atrophy in Progressive Supranuclear Palsy (PSP) and Corticobasal Syndrome (CBS) compared to healthy controls. Participants meeting diagnostic criteria for CBS or PSP, together with healthy controls, were recruited from the Cambridge Centre for Parkinson-plus. Grey matter volume, cortical thickness, and subcortical volumes were derived using FreeSurfer for 210 cortical and 36 subcortical regions defined by the Human Brainnetome Atlas. Participants with PSP were classified into Richardson syndrome (PSP-RS), corticobasal syndrome variant (PSP-CBS), frontal variant (PSP-F), and other presentations (PSP-Other). Bayesian hierarchical ANOVA models with appropriate priors were used to estimate group differences. Regions of practical equivalence (ROPE) were calculated to assess evidence of meaningful group effects. A total of 428 people were included (113 PSP-RS, 30 PSP-CBS, 23 PSP-Other, 19 PSP-F, 81 CBS, and 162 healthy controls). For grey matter volume, there was evidence of whole brain volume loss compared to controls for all disease groups except PSP-Other, which showed evidence of no difference. For cortical thickness, there was evidence of global cortical thinning in CBS compared to controls, but for no other group. For subcortical volume all groups showed evidence of volume loss compared to controls. Patterns of regional atrophy were identified in keeping with previous literature, with the largest effect observed in the left pre-motor thalamus (30% reduction in CBS). There was greater loss of subcortical volume in PSP-RS compared to other PSP subgroups. Bayesian hierarchical modelling provides a feasible framework for quantifying regional brain atrophy, with interpretable effect size estimates and uncertainty measures. This approach complements conventional statistical methods, particularly in the assessment of rare neurodegenerative and neuropsychiatric disorders.

## 1 Introduction

Identifying regions with a loss of brain volume is crucial for understanding many neurodegenerative and neuropsychiatric disorders. This is particularly relevant where regional susceptibility measured by volume loss reflects disease progression (Street et al., 2023)(Quattrone et al., 2024; Whitwell et al., 2007), or relates to change in brain activity and function (Whiteside et al., 2021).

The standard approach to assessing differences in brain structure between two disease groups utilises frequentist statistical principles to generate a *p*-value and point estimates to infer group-level differences (Ashburner & Friston, 2000; Friston, 2003; Winkler et al., 2014; Wright et al., 1995). While powerful, this approach has many limitations. For example, when there is a large sample size, a statistically “significant” result may be too small to be clinically meaningful. On the other hand, when the sample size is too small and underpowered, spurious results can be over-interpreted and significant differences missed. Discrepancies in group size may invalidate analytical approaches, forcing useful data to be discarded to balance the size of datasets. Furthermore, frequentist methods cannot be used to support evidence of similarity between groups (in favour of the null hypothesis, of no difference). Moreover, traditional statistical approaches cannot easily incorporate prior biological knowledge, for example the change in brain volume that is considered abnormal (Wagenmakers et al., 2008).

Bayesian statistics provide a potential alternative approach that addresses these limitations (Bayes, 1763; Wagenmakers et al., 2017). Bayesian approaches treat parameters as random variables and explicitly model uncertainty through probability distributions to generate an effect size and credible interval capturing the uncertainty of model parameters. They also allow the integration of prior knowledge. These advantages are especially important in rare disorders where sample sizes are often small, particularly in comparison to comparator control cohorts. Moreover, Bayesian approaches can be used to build hierarchical models which account for relationships between brain regions, since changes in brain regions are rarely independent (Lindley & Smith, 1972; Wikle, 2003).

We evaluate this approach in the context of two rare disorders: Corticobasal Syndrome (CBS) and Progressive Supranuclear Palsy (PSP). As rare and heterogeneous neurodegenerative disorders, CBS and PSP are particularly vulnerable to the limitations of frequentist approaches, and the alternative Bayesian approach may be an advantage. CBS is a heterogeneous disorder with a number of underlying neuropathologies (Alexander et al., 2014; Boeve et al., 1999; Vaughan et al., 2026), characterised by progressive asymmetric limb rigidity and bradykinesia, dystonia, myoclonus, apraxia, cortical sensory loss, and the “alien limb” phenomenon (Armstrong et al., 2013; Constantinides et al., 2019; Kouri et al., 2011). PSP classically presents with vertical supranuclear gaze palsy, early postural instability with falls, akinetic-rigid parkinsonism, poor response to levodopa, pseudobulbar palsy, and executive cognitive impairment. A variety of initial presentations are recognised which differ in their severity and prognosis (Höglinger et al., 2017; Litvan et al., 1996).

Previous studies suggest that CBS exhibits greater atrophy in cortical and basal ganglia regions than PSP, while the degree of atrophy in the midbrain and pons are similar (Dutt et al., 2016; Jabbari et al., 2020; Josephs et al., 2008). Given the clinical and imaging overlap between CBS and PSP, there is a need for improved statistical and modelling approaches to quantify regional brain changes. This motivated investigation of a Bayesian inferential framework to estimate regional brain volume loss.

The aim of this study was to assess the feasibility of evaluating regional atrophy in brain regions using a Bayesian approach: to identify differences between the rare neurodegenerative disorders PSP and CBS, and against control groups. The null hypothesis is that there is no difference between groups.

## 2 Methods

### 2.1 Patient Selection and Grouping

Through the Cambridge Centre for Parkinson-plus Disorders, we recruited participants meeting research diagnostic criteria for probable or possible CBS (Armstrong et al., 2013) or PSP (Höglinger et al., 2017), and age-matched healthy controls.

All participants underwent T1-weighted MRI imaging (slice thickness 1-1.25 mm, TE 2.93 ms, TR 2.3 s, TI 0.85 s, flip angle 8-12°). Freesurfer (version 7.2, Dale et al., 1999; Fischl et al., 1999) was used to evaluate cortical grey matter volume and average thickness of the cortex and subcortical volumes in each participant for each region of the Human Brainnetome Atlas (Fan et al., 2016) which contains 210 cortical regions and 36 subcortical regions. The PSP group were divided into subgroups depending on their initial presenting clinical features: Richardson Syndrome (PSP-RS), Corticobasal Syndrome (PSP-CBS), Frontal variant (PSP-F), and other presentations (PSP-Other). Given the likelihood of underlying PSP pathology, participants with a PSP and CBS overlap syndrome were classified under the PSP group.

### 2.2 Statistical Analyses

The distribution of volumetric measures in all comparator groups (CBS, PSP-RS, PSP-CBS, PSP-F, PSP-Other and Controls) were visualised, and a Bayesian hierarchical ANOVA model was constructed. All analysis was carried out in R (version 4.5.1) with the package brms (version 2.22.0) (Bürkner, 2017, 2018) to compare the means of cortical grey volume, average thickness of the cortex, and subcortical volume across brain regions in each diagnostic group. The predictor of interest was disease group, or subgroup. As part of a hierarchical model we estimated a common latent mean variable for all regions, and allowed both the intercept and the group effects to vary by brain region, thereby modelling region-specific differences while borrowing strength across regions through partial pooling.

These analyses were specified by the following equations:

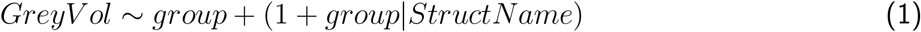

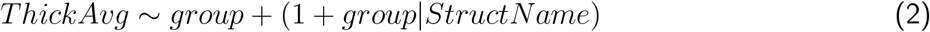

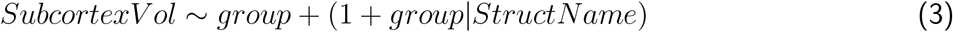

Models were estimated using the Hamiltonian Monte Carlo (HMC) Markov chain algorithm. Three chains of 6,000 iterations (including 2,000 warm-up iterations) were run, yielding 6,000 post-warmup samples. Convergence was assessed via the potential scale reduction factor 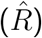 and visual inspection of trace plots.

### 2.3 Priors

We used weakly informative priors. For the fixed effects of diagnosis, we specified a Cauchy(0, 10) prior, representing the expectation that disease effects are likely centered near zero but could plausibly be large in either direction. For the intercept (corresponding to the mean value in the control group), we used a Cauchy prior centered on the observed mean of the control data, with a scale of 10, to weakly anchor the model to realistic values. For the correlations among random intercepts and slopes across brain regions, we used a Lewandowski-Kurowicka-Joe (LKJ) prior with a shape parameter of 1, which is uniform over correlation matrices. For the residual standard deviation (*σ*) and standard deviation of random effects, we used a Student-*t* prior (df = 3, *µ* = 0, sd = standard deviation of the control group), providing heavy tails while reflecting that variability is expected to be on the order of the observed standard deviation in controls, and regularising the variation allowed in intercepts and slopes across brain regions.

### 2.4 Estimating Group Differences

Group differences were assessed by examining the posterior distributions and difference between estimated group means. To make the visualisation of each brain region and its value intuitive, the effect size was defined as the percentage change of volume, i.e. the coefficient of the model divided by the standard volume of the brain region obtained from the mean of the controls.

To consider the evidence for a difference between groups, the posterior samples for each pairwise model comparison was fitted to a Bayesian linear model to obtain a Credible Interval (CI), representing the 95% distribution of the estimate for a given parameter. There are a number of methods to assess the strength of evidence for difference between groups using the posterior samples. The Bayes Factor can be used, though has limitations with smaller samples sizes, and does not reflect effect size (Sekulovski et al., 2024; Tendeiro et al., 2024). An alternative is to use the CI, for example to assess whether there is any overlap between the estimates of the probability distribution. However, trivial effect sizes can still be considered important using this approach. Therefore we used a Region Of Practical Equivalence (ROPE) that represents a distribution within which there is no meaningful difference between the groups (Kruschke, 2018). We defined the ROPE to be 5% change from the control group mean.

## 3 Results

We analysed data from 428 participants (113 PSP-RS, 30 PSP-CBS, 23 PSP-Other, 19 PSP-F, 81 CBS, 162 healthy controls). Table 1 shows the demographics of the patients in each group.

**Table 1:** Demographic Characteristics by Group. Values are shown as mean ± standard deviation for age, years of education, ACE-R, PSPRS and UPDRS. CBS = Corticobasal syndrome; PSP-CBS = Progressive Supranuclear Palsy, Corticobasal Syndrome variant; PSP-F = Progressive Supranuclear Palsy, Frontal variant; PSP-Other = Progressive Supranuclear Palsy, other variants; PSP-RS = Progressive Supranuclear Palsy, Richardson Syndrome.

| Group | PSP-RS | PSP-CBS | PSP-F | PSP-Other | CBS | Control |
| --- | --- | --- | --- | --- | --- | --- |
| Sex (M/F) | 55/58 | 14/16 | 13/6 | 13/10 | 39/42 | 85/77 |
| Age | 71.1 ± 6.8 | 68.3 ± 7.3 | 68.1 ± 8.2 | 72.2 ± 5.9 | 69.1 ± 8.4 | 66.8 ± 9.1 |
| Years of Education | 14.2 ± 4.0 | 14.0 ± 4.6 | 13.0 ± 4.2 | 13.5 ± 3.5 | 12.9 ± 2.9 | 15.2 ± 4.0 |
| ACE-R | 75.3 ± 15.9 | 79.0 ± 11.5 | 76.7 ± 17.6 | 84.9 ± 8.6 | 69.6 ± 21.5 | 92.9 ± 8.8 |
| PSPRS | 31.9 ± 13.1 | 30.7 ± 12.2 | 31.0 ± 14.7 | 22.8 ± 11.7 | 30.7 ± 16.9 | - |
| UPDRS | 28.5 ± 14.8 | 29.9 ± 10.2 | 29.1 ± 18.3 | 24.5 ± 13.3 | 31.5 ± 18.8 | - |

Average group differences across all regions (i.e. the hierarchical model latent variable for regional differences) for cortical volume, thickness, and subcortical volume are summarised in Table 2. For each measure the CBS group was associated with the greatest decrease with reference to the control group. The 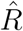 values for all the parameters was less than 1.05, meaning that the models had converged.

**Table 2:** Percentage change differences with 95% CI between the control group and disease groups in grey matter volume, average thickness and subcortical volume. All groups showed evidence of difference from the control group. CBS = Corticobasal syndrome; PSP-CBS = Progressive Supranuclear Palsy, Corticobasal Syndrome variant; PSP-F = Progressive Supranuclear Palsy, Frontal variant; PSP-Other = Progressive Supranuclear Palsy, other variants; PSP-RS = Progressive Supranuclear Palsy, Richardson Syndrome.

| Disease Group | Grey Matter Volume | Average Cortical Thickness | Subcortical Volume |
| --- | --- | --- | --- |
| CBS | -14.0% (-15.0%; -12.9%) | -5.74% (-6.56%; -5.33%) | -15.6% (-17.5%; -13.6%) |
| PSP-CBS | -7.50% (-8.26%; -6.68%) | -2.46% (-2.87%; -2.46%) | -9.86% (-11.9%; -7.71%) |
| PSP-F | -9.85% (-10.9%; -8.80%) | -2.87% (-3.69%; -2.46%) | -15.0% (-17.6%; -12.3%) |
| PSP-RS | -11.4% (-12.1%; -10.5%) | -2.46% (-2.87%; -2.46%) | -16.6% (-19.0%; -14.2%) |
| PSP-Other | -2.86% (-3.63%; -2.08%) | -1.64% (-2.05%; -1.23%) | -7.09% (-8.97%; -5.11%) |

Figures 1,2, and 3 show the group differences and ROPE for the parameters of the average cortical thickness, average cortical thickness and subcortical volume against healthy controls. The overlap between the credible interval (CI) estimate and ROPE of all parameters for grey matter volume was 0% providing strong evidence of group difference, except for PSP-Other, where the overlap ROPE was 100% providing strong evidence in favour of the null hypothesis (i.e. no difference between groups). However, for cortical thickness, all disease groups, except CBS, had an overlap between the CI and ROPE of 100% providing strong evidence for the null hypothesis. For subcortical volume, the overlap between the CI and ROPE was 0% for all disease groups except the PSP-Other group that had i5% overlap, providing strong evidence of differences between all groups and healthy controls.

**Figure 1.**
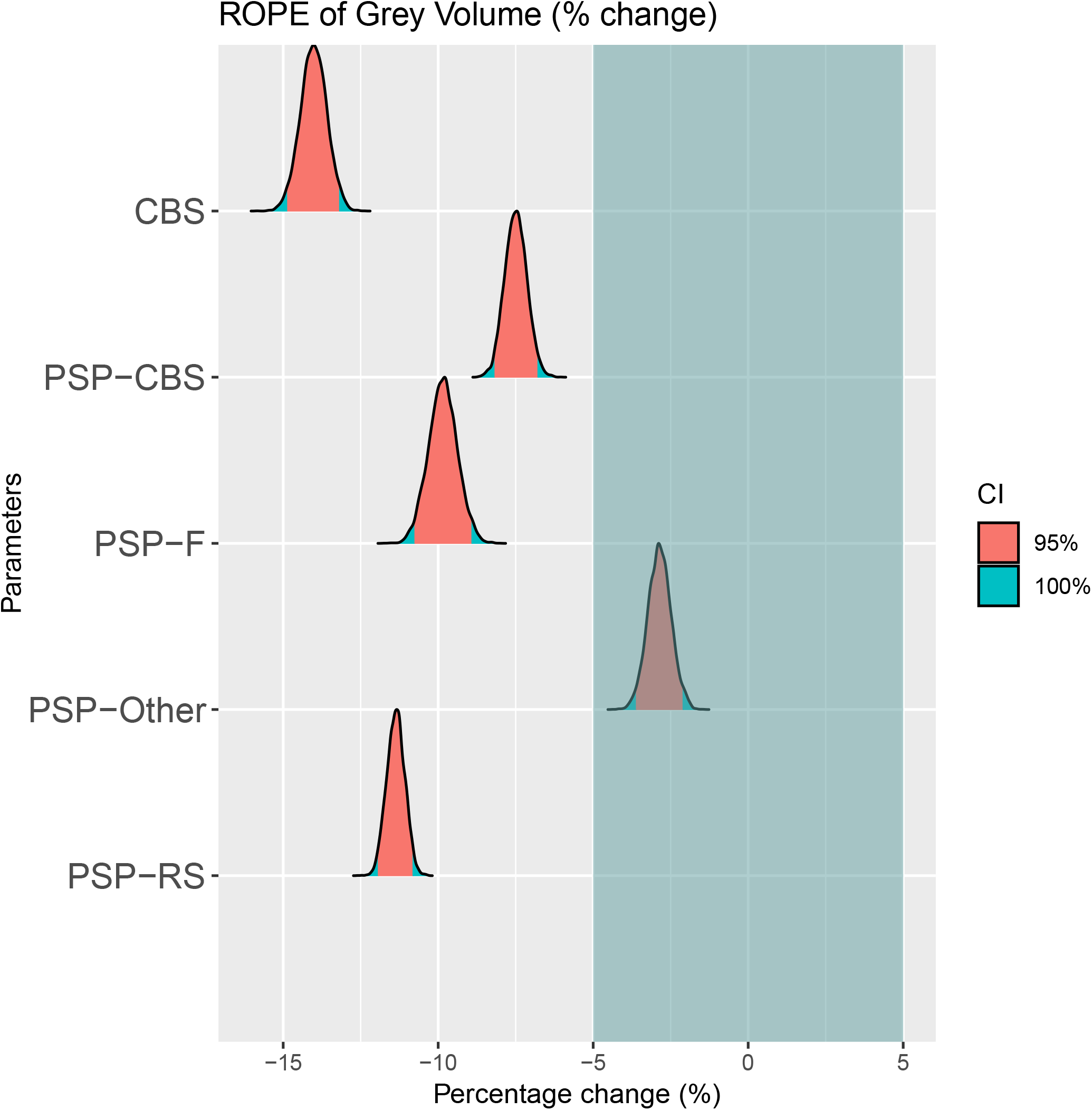
Grey matter volume posterior distributions of all diagnostic groups, showing the Region of Practical Equivalence (ROPE) shaded in green. CBS = Corticobasal syndrome; PSP-CBS = Progressive Supranuclear Palsy, Corticobasal Syndrome variant; PSP-F = Progressive Supranuclear Palsy, Frontal variant; PSP-Other = Progressive Supranuclear Palsy, other variants; PSP-RS = Progressive Supranuclear Palsy, Richardson Syndrome.

**Figure 2.**
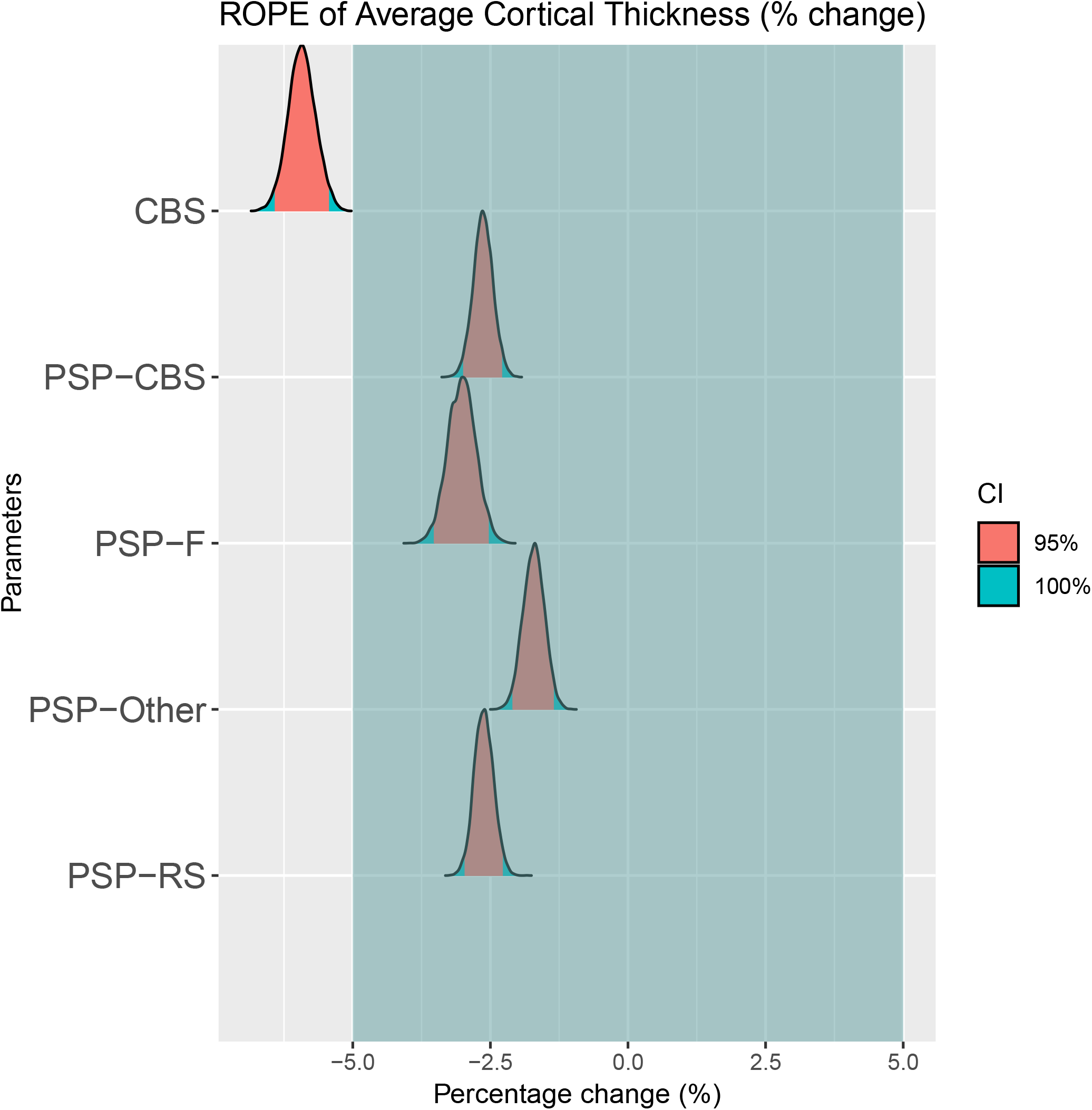
Cortical thickness posterior distributions of all diagnostic groups, showing the Region of Practical Equivalence (ROPE) shaded in green. CBS was the only group demonstrating evidence for a difference. All other groups showed a difference smaller than the ROPE, suggesting no evidence for a difference between groups. CBS = Corticobasal syndrome; PSP-CBS = Progressive Supranuclear Palsy, Corticobasal Syndrome variant; PSP-F = Progressive Supranuclear Palsy, Frontal variant; PSP-Other = Progressive Supranuclear Palsy, other variants; PSP-RS = Progressive Supranuclear Palsy, Richardson Syndrome.

**Figure 3.**
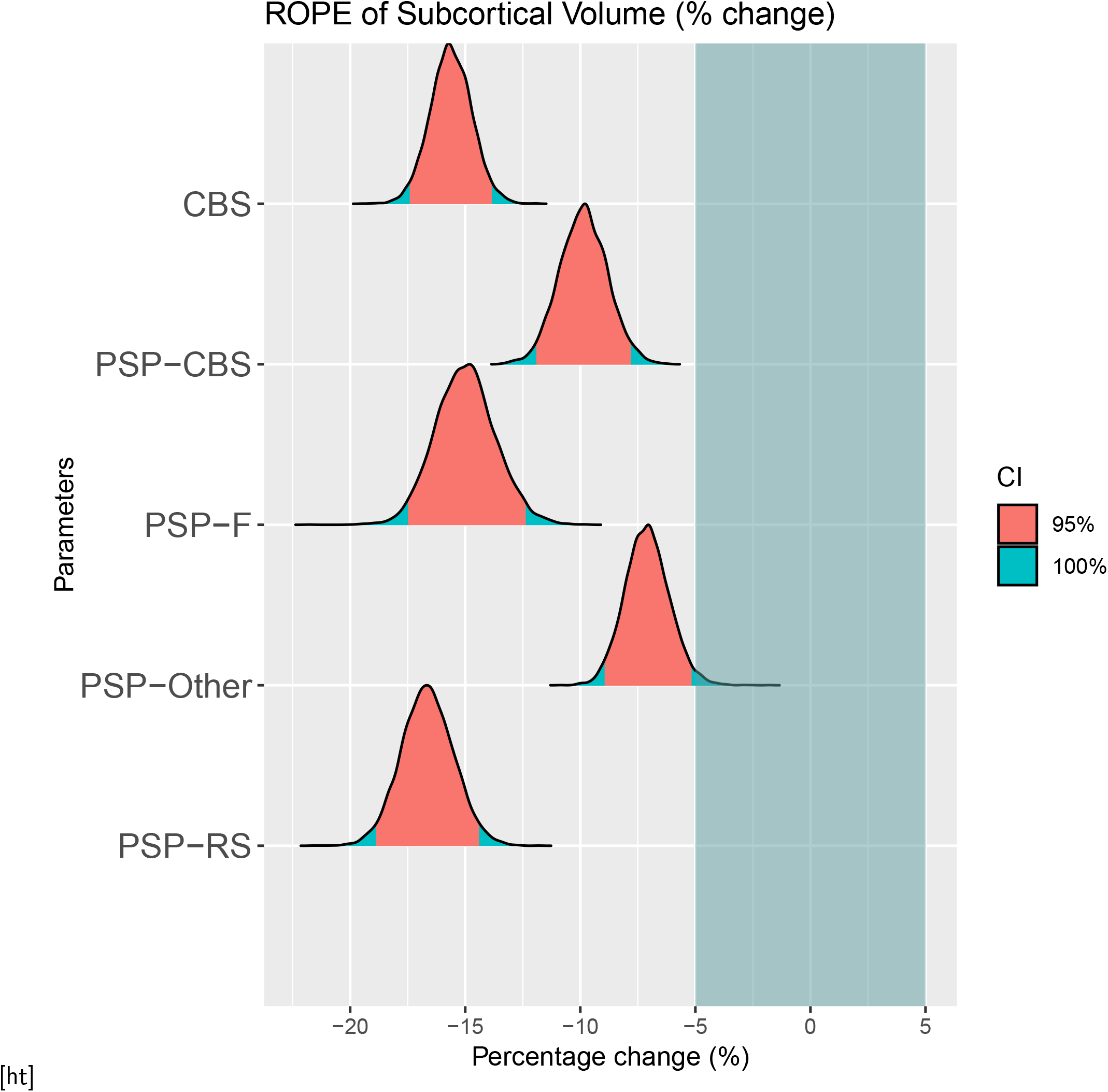
Subcortical volume posterior distributions of all diagnostic groups, showing the Region of Practical Equivalence (ROPE) shaded in green. CBS = Corticobasal syndrome; PSP-CBS = Progressive Supranuclear Palsy, Corticobasal Syndrome variant; PSP-F = Progressive Supranuclear Palsy, Frontal variant; PSP-Other = Progressive Supranuclear Palsy, other variants; PSP-RS = Progressive Supranuclear Palsy, Richardson Syndrome.

To visualise the grey matter volume, cortical thickness, and subcortical volume in specific brain regions, we plotted the percentage volume change by region for each disease group (Figures 4, 5 and 6). The brain regions in the CBS case showed the greatest change. The top 5 most affected brain regions for each disease and for each metric (cortical grey volume, average cortical thickness and subcortical volume) are summarised in Table 4.

**Table 3:** Pairwise comparison for grey matter volume, average cortical thickness and subcortical volume for different disease groups. CBS = Corticobasal syndrome; PSP-CBS = Progressive Supranuclear Palsy, Corticobasal Syndrome variant; PSP-F = Progressive Supranuclear Palsy, Frontal variant; PSP-Other = Progressive Supranuclear Palsy, other variants; PSP-RS = Progressive Supranuclear Palsy, Richardson Syndrome.

| Hypothesis | Grey Matter Volume (mm<sup>3</sup>) | Average Cortical Thickness (mm) | Subcortical Volume (mm<sup>3</sup>) |
| --- | --- | --- | --- |
| CBS vs PSP-CBS | -142.72 (-159.90; -125.66 ) | -0.08 (-0.09; -0.07) | -78.55 (-100.83; -56.47) |
| CBS vs PSP-RS | -57.98 (-70.31; -45.58) | -0.08 (-0.09; -0.07) | 13.97 (-7.50; 34.81) |
| CBS vs PSP-F | -91.23 (-108.62; -73.41) | -0.07 (-0.08; -0.06) | -8.58 (-38.35; 20.11) |
| CBS vs PSP-Other | -244.11 (-263.40; -223.09) | -0.10 (-0.11; -0.09) | -116.37 (-141.19; -91.74) |
| PSP-CBS vs PSP-RS | 84.74 (71.51; 98.24) | 0.00 (-0.01; 0.01) | 92.52 (70.69; 113.82) |
| PSP-CBS vs PSP-F | 51.49 (32.61; 70.17) | 0.01 (0.00; 0.02) | 69.97 (41.78; 98.41) |
| PSP-CBS vs PSP-Other | -101.39 (-118.97; -82.78) | -0.02 (-0.03; -0.01) | -37.82 (-65.49; -9.26) |
| PSP-RS vs PSP-F | -33.26 (-49.43; -17.49) | 0.01 (0.00; 0.02) | -22.55 (-45.11; 0.62) |
| PSP-RS vs PSP-Other | -186.13 (-202.58; -168.14) | -0.02 (-0.03; -0.01) | -130.35 (-158.45; -102.27) |
| PSP-F vs PSP-Other | -152.88 (-173.60; -131.45) | -0.03 (-0.04; -0.02) | -107.79 (-140.19; -74.16) |

**Table 4:**
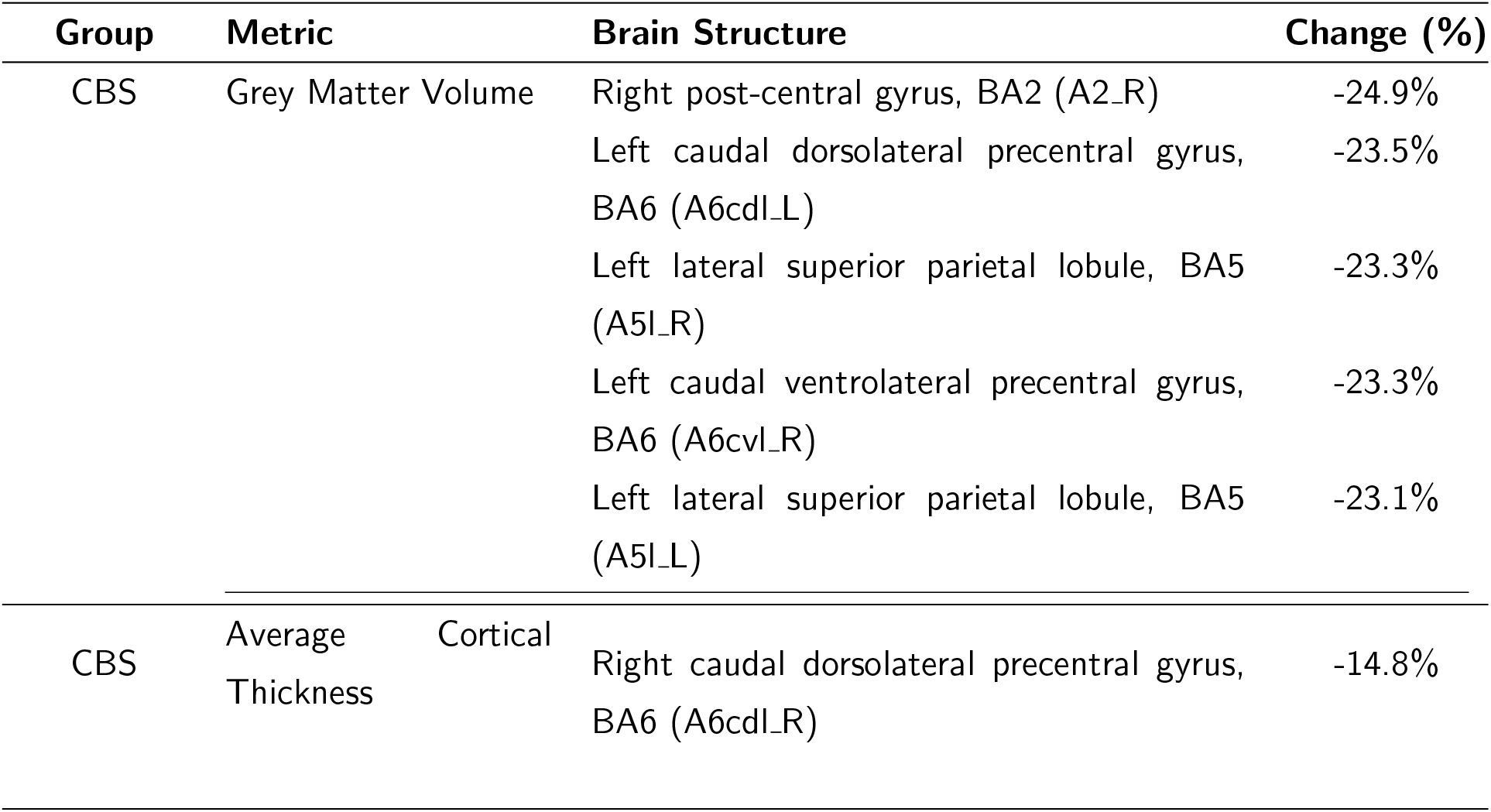
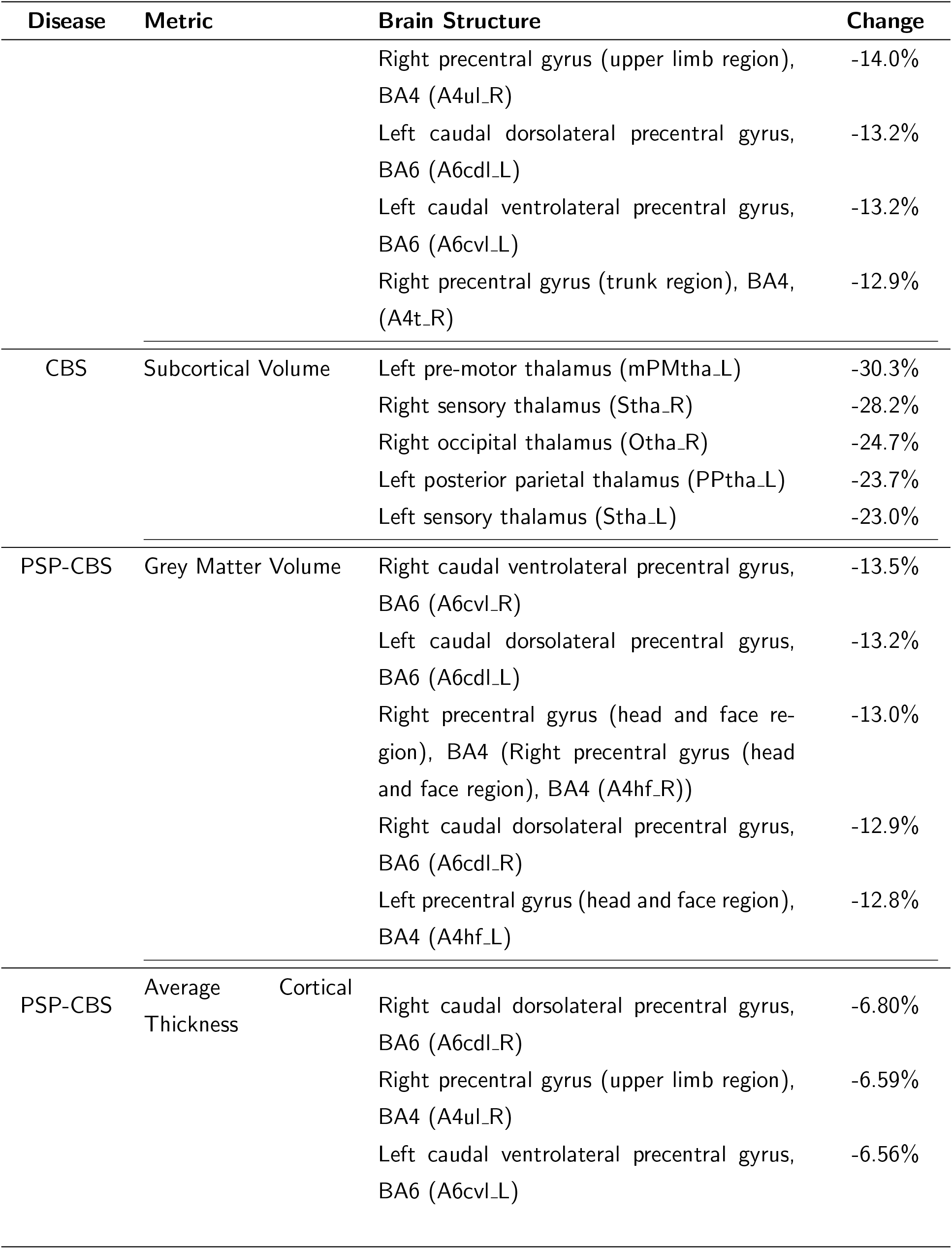
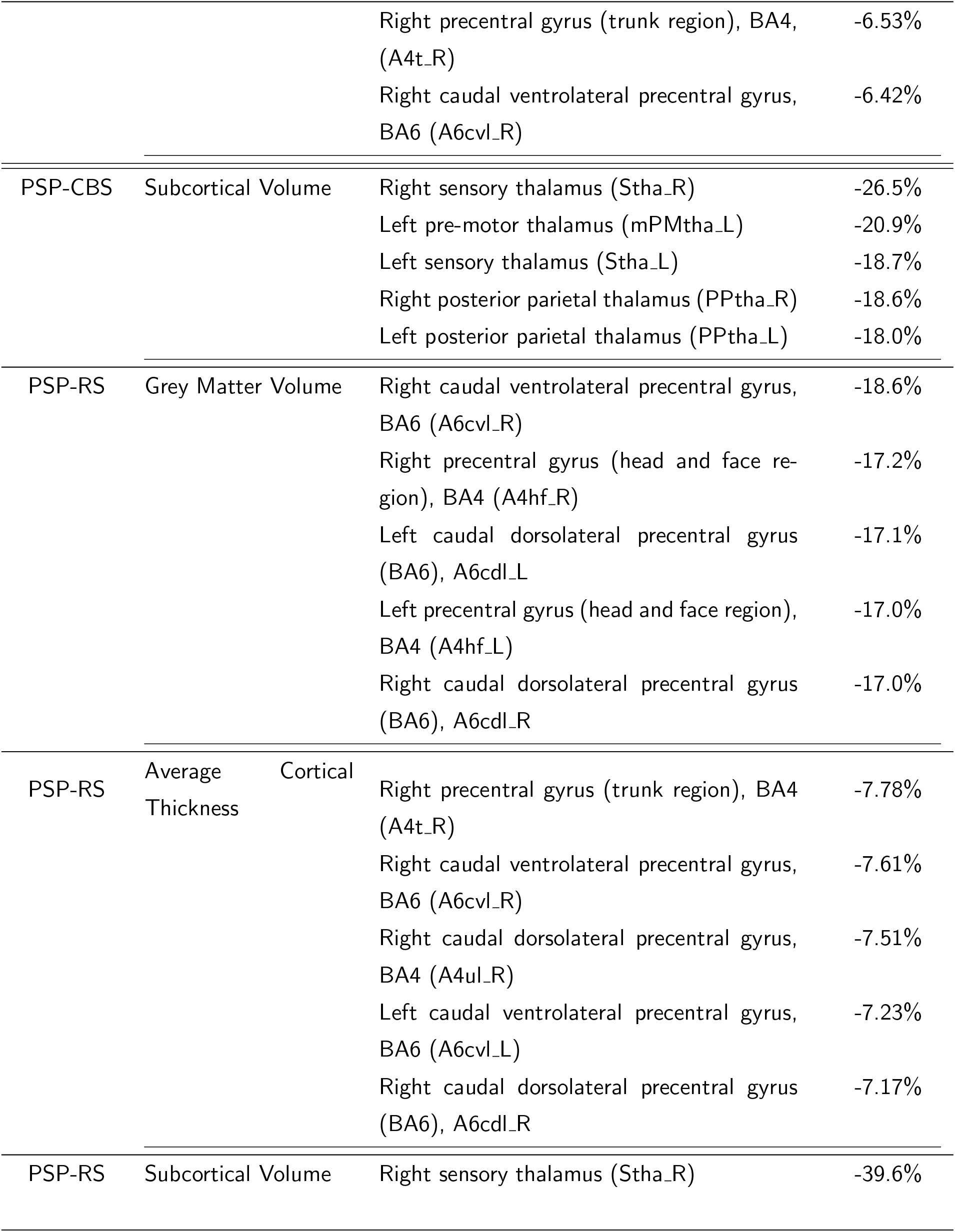
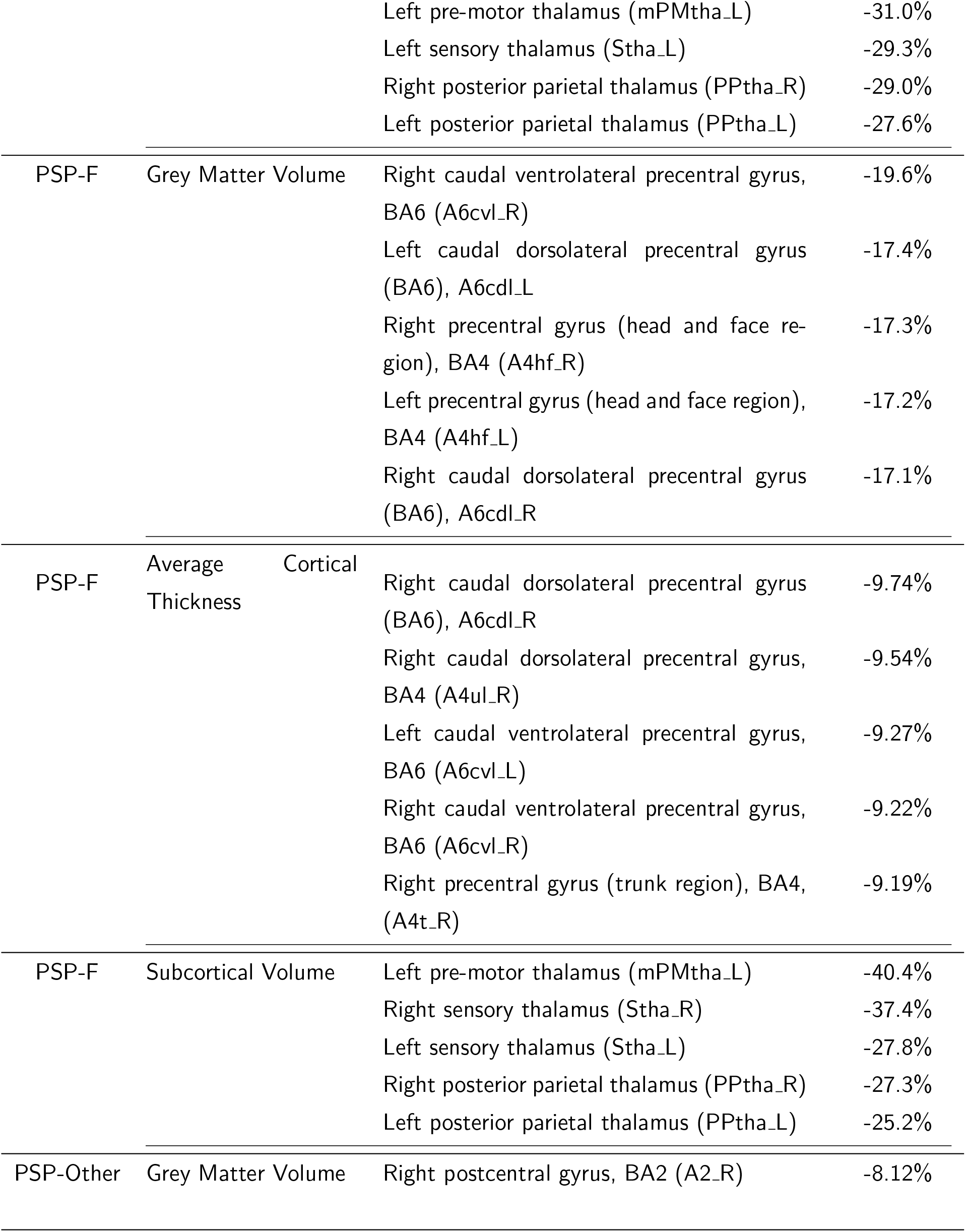
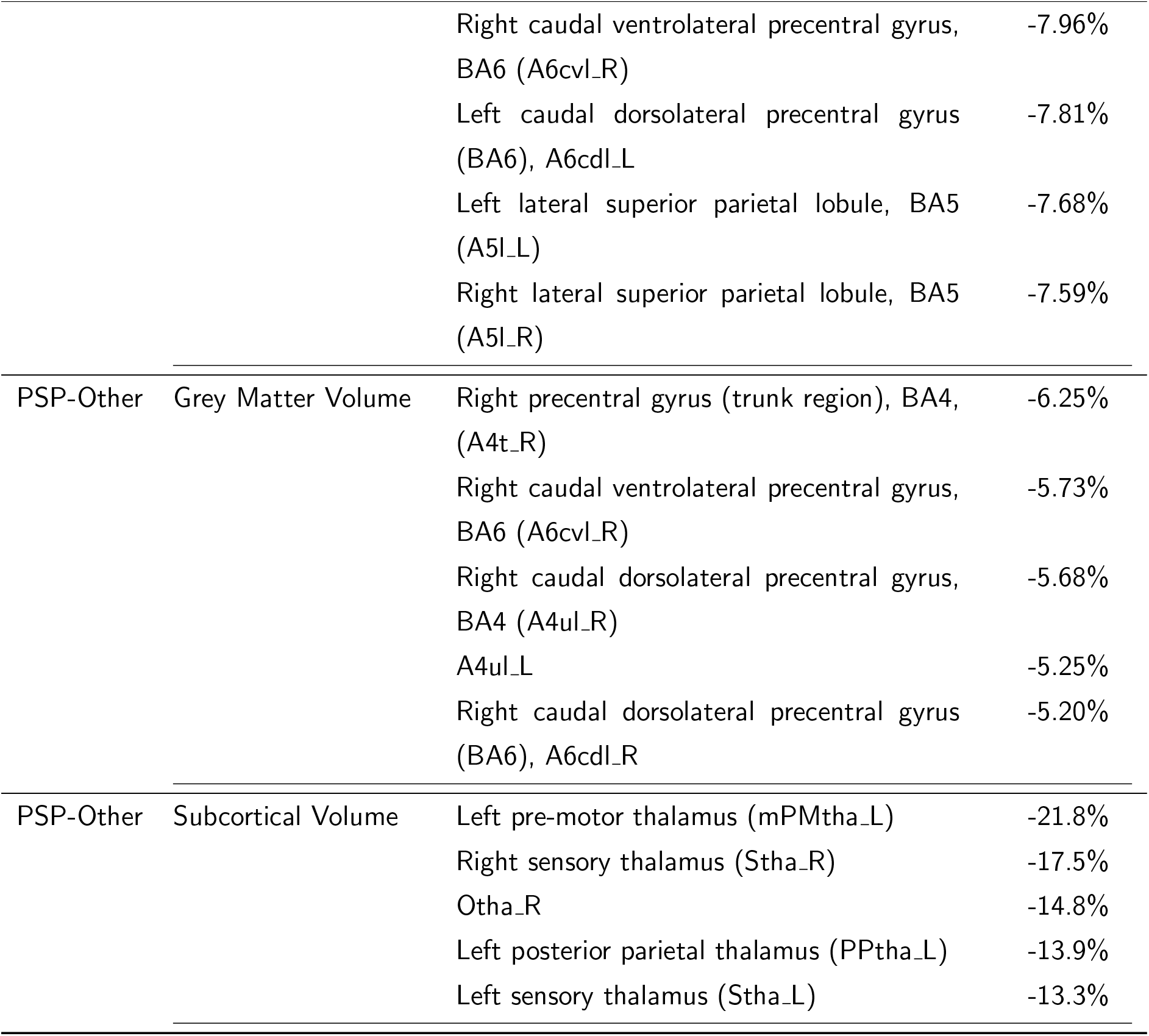
Top 5 brain regions for each disease group for cortical grey matter volume, mean cortical thickness and subcortical volume. CBS = Corticobasal syndrome; PSP-CBS = Progressive Supranuclear Palsy, Corticobasal Syndrome variant; PSP-F = Progressive Supranuclear Palsy, Frontal variant; PSP-Other = Progressive Supranuclear Palsy, other variants; PSP-RS = Progressive Supranuclear Palsy, Richardson Syndrome.

**Figure 4.**
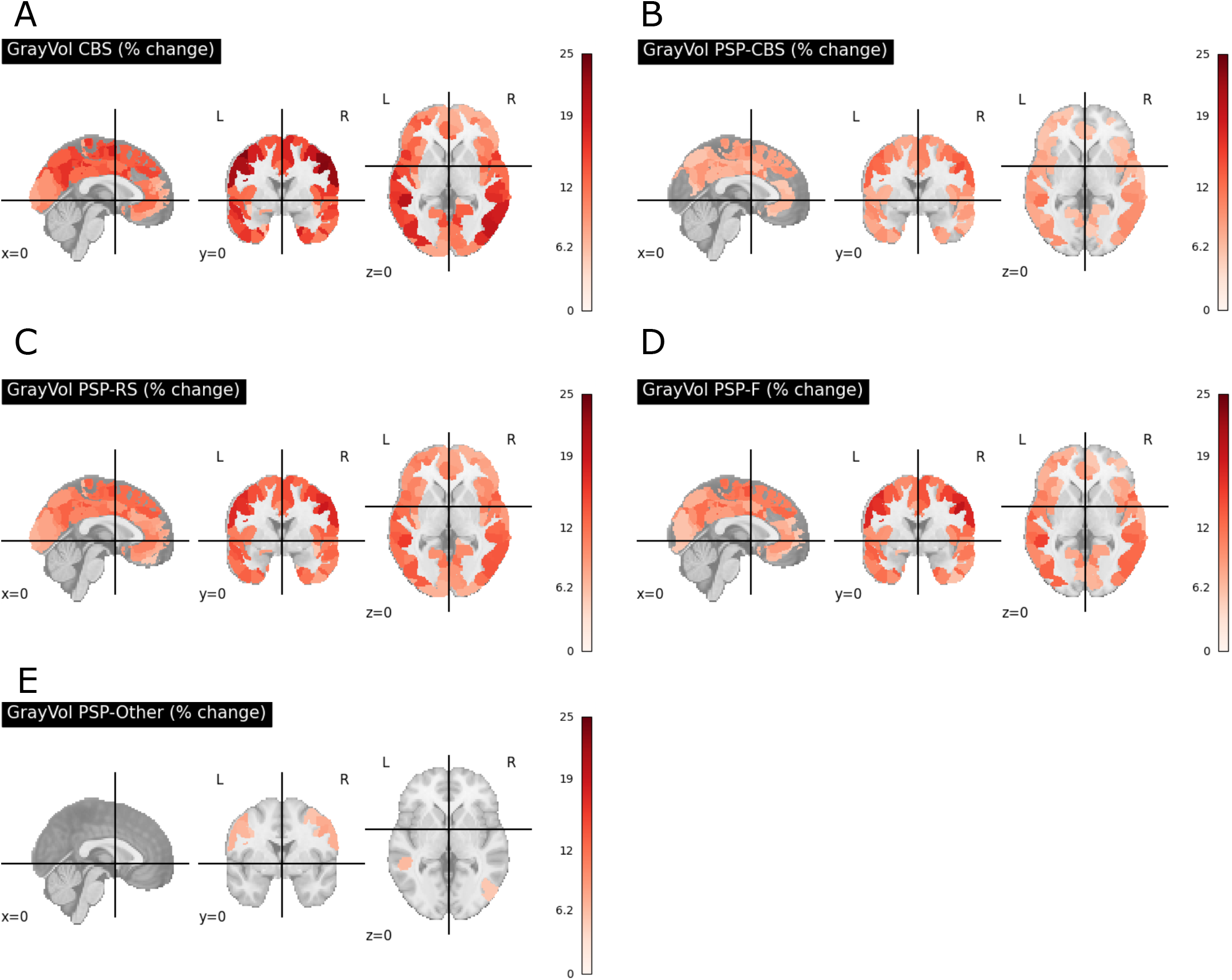
Percentage change of the grey volume of brain cortical areas for A: CBS, B: PSP-CBS, C: PSP-RS, D: PSP-F, E: PSP-Other CBS had the greatest percentage loss of cortical grey matter volume. CBS = Corticobasal syndrome; PSP-CBS = Progressive Supranuclear Palsy, Corticobasal Syndrome variant; PSP-F = Progressive Supranuclear Palsy, Frontal variant; PSP-Other = Progressive Supranuclear Palsy, other variants; PSP-RS = Progressive Supranuclear Palsy, Richardson Syndrome.

**Figure 5.**
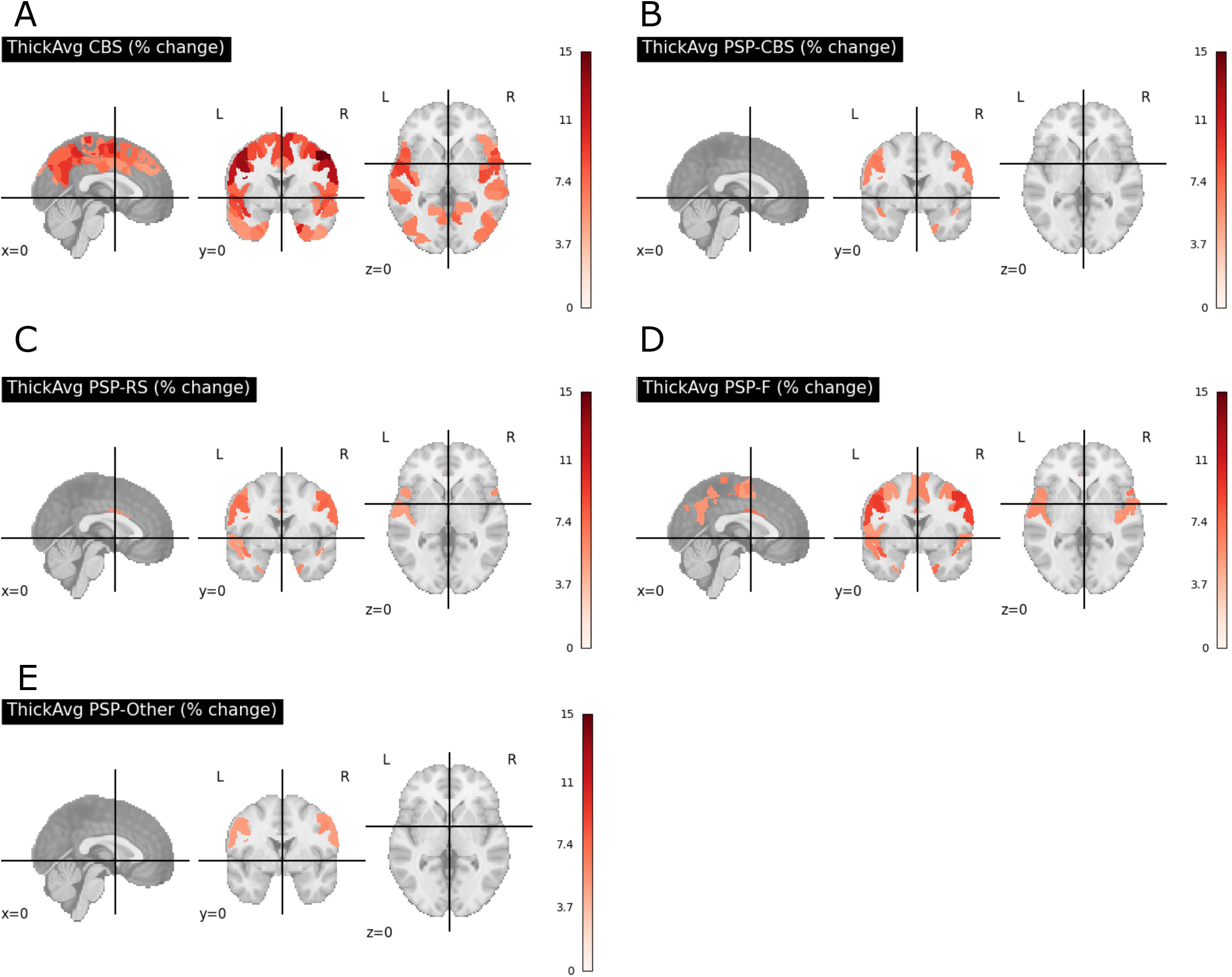
Percentage change of the average thickness of brain cortical areas for A: CBS, B: PSP-CBS, C: PSP-RS, D: PSP-F, E: PSP-Other CBS has the greatest percentage loss of cortical thickness. CBS = Corticobasal syndrome; PSP-CBS = Progressive Supranuclear Palsy, Corticobasal Syndrome variant; PSP-F = Progressive Supranuclear Palsy, Frontal variant; PSP-Other = Progressive Supranuclear Palsy, other variants; PSP-RS = Progressive Supranuclear Palsy, Richardson Syndrome.

**Figure 6.**
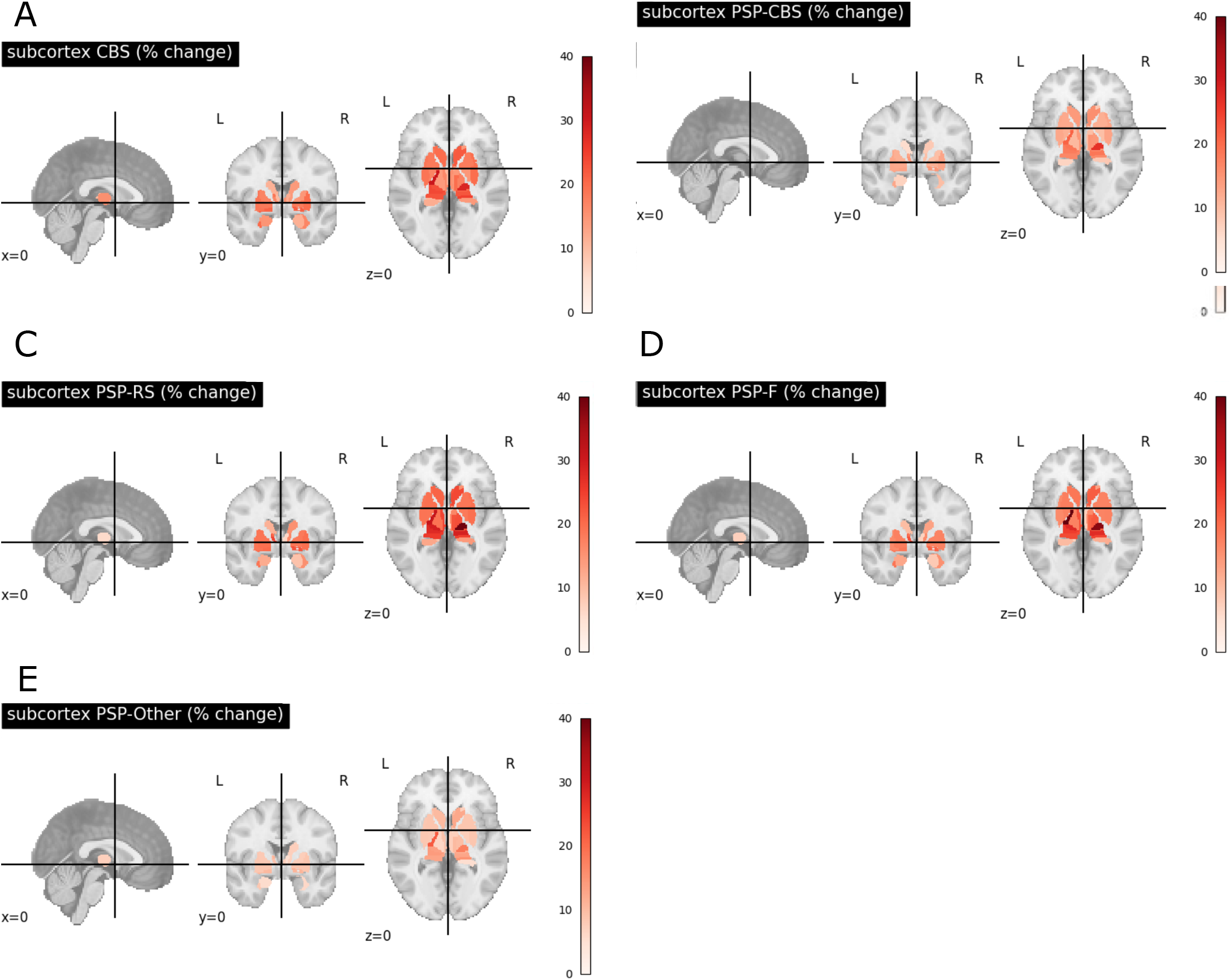
Percentage change of the average thickness of brain cortical areas for A: CBS, B: PSP-CBS, C: PSP-RS, D: PSP-F, E: PSP-Other CBS has the greatest percentage loss of subcortical volume. CBS = Corticobasal syndrome; PSP-CBS = Progressive Supranuclear Palsy, Corticobasal Syndrome variant; PSP-F = Progressive Supranuclear Palsy, Frontal variant; PSP-Other = Progressive Supranuclear Palsy, other variants; PSP-RS = Progressive Supranuclear Palsy, Richardson Syndrome.

Differences between disease groups are shown in Table 3. For grey matter volume, there were non-overlapping CIs between all disease groups, suggesting at least modest evidence of difference between all groups. Given this analysis examined difference between groups rather than comparing to controls, we did not consider the ROPE. For cortical thickness, there was no evidence of difference between group differences in the average thickness between PSP-CBS and PSP-RS, between PSP-CBS and PSP-F, or betweeen PSP-RS and PSP-F. For the subcortical volume, there was no evidence of difference between the CBS and PSP-RS group, or between the CBS and PSP-F group.

## 4 Discussion

This study demonstrates the feasibility and utility of Bayesian statistical analysis to identify meaningful and clinically relevant structural change in rare neurodegenerative disorders. Specifically, we have identified robust differences in grey matter volume and cortical thickness between control, CBS, and PSP subtypes using a Bayesian hierarchical modeling approach. Similar brain regions demonstrated atrophy across CBS, PSP-CBS, PSP-RS and PSP-F. In keeping with previous literature, we found evidence that CBS has a greater extent and degree of cortical atrophy than the other diagnostic groups. We also find that people with the Richardson syndrome of PSP had evidence of greater subcortical volume loss than either PSP-CBS or other presentations.

The difference between PSP-RS and PSP-CBS/PSP-Other groups likely reflects the localisation, stage and progression of PSP neuropathology. Subtypes of PSP typically present with a specific syndrome before typical PSP-RS clinical features emerge some years into the disease course (Höglinger et al., 2017; Respondek et al., 2014). The PSP-Other group in particular encompasses syndromes recognised to be more slowly progressive, such as Primary Gait Freezing (Compta et al., 2007; Williams et al., 2007), the ocular variant of PSP (Respondek et al., 2014), and PSP-parkinsonism (Williams et al., 2005) which is an akinetic-rigid syndrome distinguished from PSP-RS by a later onset of falls.

Confidence and reproducibility of results can be challenging in neuroimaging studies, particularly where groups are small or differ in size, and there is complex structure within the data. Our approach provides mitigation against these concerns. Firstly, the Bayesian approach using Regions of Practical Equivalence (ROPEs) allow the inclusion of clinical knowledge to provide informed cut-offs of effect size (Kruschke, 2018); in our case 5% change in regional volume or thickness. We accept that the 5% threshold is somewhat arbitrary; it is perhaps a conservative estimate of a difference which is meaningful. Future studies may explore whether meaningful thresholds of difference vary between regions and between disorders.

Secondly, the precision of model parameters can be estimated in Bayesian models, and depends on the size and distribution of the observed dataset reflected in posterior distributions (Zitzmann & Hecht, 2019). This is particularly valuable in small groups and complex data structures, where frequentist approaches may be limited by degrees of freedom, or fail to account for differences in distribution between groups (Baldwin & Fellingham, 2013). Using weakly informative priors, each group can be modelled independently with little expectation of the final distribution.

Finally, using a hierarchical model allows specific regional differences whilst properly accounting for the non-independence between regions by partially pooling between group differences. The non-independence in frequentist approaches is often addressed by permissive correction for multiple comparisons (e.g. using the False Discovery Rate (Benjamini & Yekutieli, 2001)), or using clustering techniques (e.g. Threshold-Free Cluster Enhancement (Smith & Nichols, 2009)) that rely on adjustment of *p*-value estimates. Multiple comparison methods can still be overly conservative when a large number of regions are assessed. TFCE and similar methods are usually used for voxel-wise analysis where they are more computationally efficient than Bayesian hierarchical approaches, but the significance of the difference is determined by cluster size which might lead to over-estimation of differences between adjacent regions, and may overlook smaller regions of difference.

Given their advantages, hierarchical Bayesian models have been applied to functional MRI and other imaging modalities (Friston, Penny, et al., 2002). In this context, parametric empirical Bayes (PEB) estimators were used as an alternative to maximum likelihood estimates for a second level group analysis accounting for individual differences (Friston, Glaser, et al., 2002). Although the PEB approach has been applied to joint inference using structural and functional connectivity MRI (Sokolov et al., 2019), we have not previously seen these methods applied to more traditional volumetric neuroimaging analysis.

There are several limitations to our study. Firstly, the data that we obtained are from a single centre in the UK which might not generalise to other centres in the UK or across the world. Secondly, as PSP and CBS are rare diseases, the number of cases are very low for some subgroups. Thirdly, the hierarchical model, whilst allowing information sharing between brain regions, is very computationally expensive and requires the use of high performance computing to run. This makes it very difficult to scale the approach to a large amount of data. However, cohorts of rare disorders typically have numbers not exceeding hundreds, which are computationally tractable for most grid computing infrastructures.

## 5 Conclusion

In this study, we use a Bayesian hierarchical model to examine structural brain changes in PSP and CBS compared to controls. We demonstrate the model’s utility in finding differences in brain volume between diagnostic groups and subgroups. We show that Bayesian hierarchical models improve the confidence in estimating structural brain differences in the context of rare neurological disorders.

## 6 Declaration on the use of AI

We did not use AI in writing this paper.

## 7 Data and Code Availability

Data is available upon reasonable request. Code for the project is available at: gitlab.developers.cam.ac.uk/tr332/bayes structural

## 8 Author contributions

TR and SYSH were involved in conceptualisation, data curation, formal analysis, investigation, methodology, software and writing the original draft. TS was involved in methodology, software, and reviewing and editing the writing. BG AND JBR were involved in data curation, and reviewing and editing the writing.

## 11 Acknowledgments

We are grateful to our wonderful patients and their families who support our research.

